# CLEAR-FORCE: A reservoir computing framework for closed-loop brain stimulation

**DOI:** 10.64898/2026.10.05.756709

**Authors:** Athena Moreno-Gris, Camille Godin, Jean-Philippe Thivierge

## Abstract

Electrical stimulation of brain activity has emerged as a key therapeutic intervention for neurological diseases. The success of these approaches relies on precisely tuned stimulation parameters. A major challenge is that these parameters should ideally be adjusted in real-time based on changes in brain activity. Closed-loop controllers address this challenge by monitoring brain activity and adapting stimulation according to ongoing changes in neural dynamics. While machine learning has been helpful in this context, closed-loop controllers typically do not adapt in real-time to fluctuations in brain activity. Here, we propose a novel framework based on reservoir computing to train a closed-loop, real-time neural controller. This approach, termed CLEAR-FORCE, trains the output weights of a recurrent network based on a combination of target signals and feedback from brain activity. Two series of simulations were performed, where the controller was successful at (1) inducing neural oscillations in a population of spiking neurons and (2) suppressing pathological oscillations. This framework provides a promising avenue for the implementation of neural controllers in low-powered, real-time, and adaptive hardware that harnesses machine learning for the personalized treatment of brain disorders.

## I. Introduction

In recent years, neuromodulation of brain activity has become a key form of therapy for neurological and psychiatric conditions that do not respond to conventional treatments [1-3]. By delivering mild electrical stimulation to specific brain regions, symptoms can be alleviated in disorders such as Parkinson’s disease, Tourette syndrome, and essential tremor [4-6]. Available methods for delivering electrical activity include deep brain stimulation, transcranial electrical stimulation, and transcranial alternating current stimulation [7].

Although clinically successful in many cases, existing stimulation approaches depend on precisely tuned stimulation parameters such as the shape of the oscillatory wave, its amplitude, and its frequency [8]. These parameters are manually adjusted by clinicians, and recalibration is often required as the patient’s brain state changes over time. Because the parameter space is high-dimensional and patient-specific, this process can be time-consuming and inefficient [9]. Moreover, most devices operate in an open-loop mode, where stimulation is delivered without regard to real-time neural activity. This makes it difficult to adapt therapy to fluctuations in symptoms or brain dynamics that occur on timescales ranging from minutes to weeks and months.

Closed-loop stimulation aims to address these shortcomings by continuously monitoring neural activity and adapting stimulation parameters in real time [10]. Landmark studies have demonstrated the benefits of this approach. For example, adaptive deep brain stimulation improved motor outcomes in Parkinson’s patients compared with constant stimulation [11]. Similar strategies have shown success in epilepsy, where seizure-related patterns in intracranial EEG are used to trigger preventative stimulation [12].

To guide closed-loop stimulation, recent years have seen a growing interest in machine learning (ML) approaches for identifying patient-specific stimulation strategies [13,14]. Several methods have been tested in both simulations and clinical settings [15-18]. For instance, recent efforts have designed a controller based on Bayesian optimization [19]. Other approaches have employed deep neural networks to predict neural responses to stimulation [20-23]. Personalized brain decoders using ML are emerging for disorders such as epilepsy, depression, and movement impairments [24-28]. Implementations of ML in hardware, such as system-on-chips for seizure control, further underscore the promise of these approaches in neuromodulation [29-33].

Many ML approaches, however, do not operate in real time and have not been validated for both enhancing and suppressing brain oscillations. The complexity of brain activity presents a challenge to developing these approaches. Neural systems exhibit “organized complexity”, where outcomes depend not only on the properties of individual elements but also on their nonlinear interactions [7]. This makes it difficult to predict how stimulation will influence neural dynamics over time. A computational framework that captures and forecasts such complex activity may therefore play a central role in improving neuromodulation.

One class of neural networks that has shown promises for the control of neural activity is reservoir computing [34,35]. In this approach, a time-evolving recurrent neural network projects its activity to readout nodes via trainable weights that are adjusted in real time to minimize a desired error function. This approach reproduces chaotic and nonlinear brain dynamics and can implement a wide variety of time-dependent signals from simple oscillations to complex patterns of activity.

In this work, we introduce a novel closed-loop framework where a reservoir network termed CLEAR-FORCE (Closed-Loop Error-based Adaptive Regulation, based on First-Order, Reduced and Controlled Error [35]) is employed as a neural controller capable of both synchronizing and desynchronizing spiking neurons. We show that this model can adapt neural stimulation to meet specific therapeutic goals. Going further, we discuss the potential implementation of reservoir models in low-power neural stimulation hardware [10].

The remainder of this work is structured as follows. Section II describes the reservoir controller and its application to regulating simulated brain activity at the level of spiking neurons. Section III demonstrates the use of this framework to induce neural oscillations. Section IV examines how the neural controller may be employed to desynchronize pathological brain activity. Section V provides estimates of power consumption for neuromorphic implementation. Finally, the Discussion highlights pathways toward clinical applications.

## II. CLEAR-FORCE CONTROLLER AND SPIKING NETWORK DYNAMICS

The proposed framework involves two main components (Fig. 1). First, a neural controller consists of a reservoir network that controls the output delivered to a target brain region and receives feedback from that region in order to adjust its output. Second, the target brain region is composed of randomly interconnected spiking neurons that receive electrical activity from the controller.

**Fig. 1.**
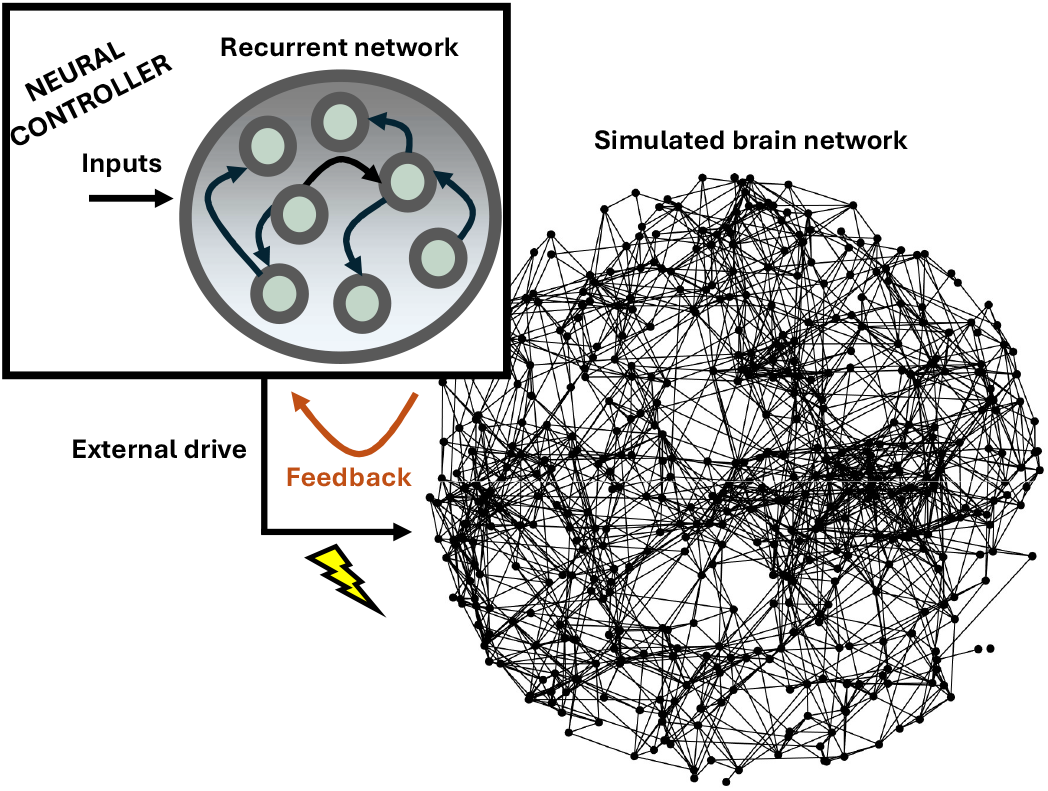
Schema of the neural controller combined with a simulated brain network that receives neural stimulation and sends feedback to the controller in a closed-loop fashion.

Together, these two components form a closed loop where feedback from brain activity influences the controller. Importantly, the reservoir learns in real time while receiving ongoing feedback from the spiking network. This approach differs from systems where the training phase is distinct from the stimulation phase [36].

The proposed closed loop pipeline is useful in clinical contexts for two main reasons. First, the impact of neural stimulation on brain activity is largely unknown, as the complexity of brain dynamics prevents a straightforward mapping between stimulation parameters and neural responses. Second, brain dynamics are non-stationary [7], hence an adaptive closed-loop controller is needed to track changes in activity that evolve over time.

Next, we provide an overview of the two components in turn, namely the simulated brain network and the neural controller. A full processing pipeline is presented in Fig. 2 and complete open-source code is available from https://doi.org/10.6084/m9.figshare.31158112.

**Fig. 2.**
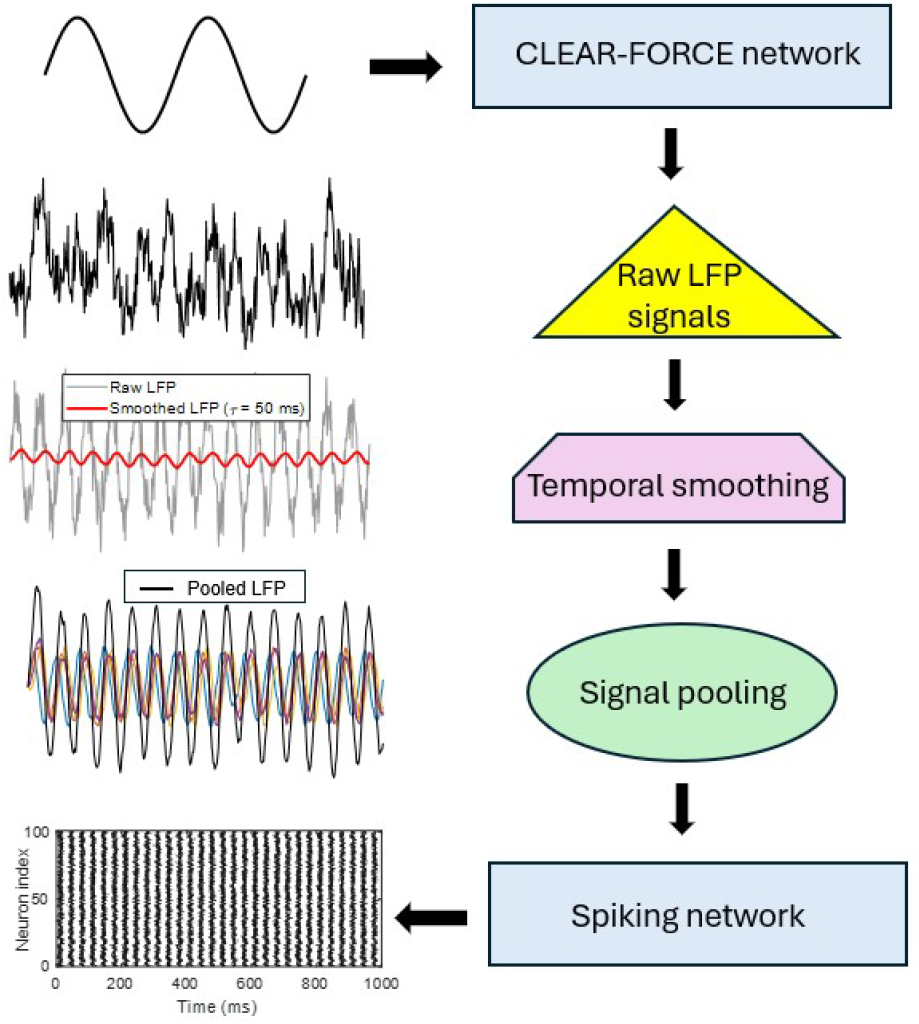
Processing pipeline for the neural controller combined with CLEAR-FORCE learning. For ease of visualization, the arrow looping from the spiking network back to the CLEAR-FORCE network was omitted.

### A. Simulated brain network

The brain network targeted by the neural controller was simulated as a population of interconnected *N*=100 spiking neurons (unless otherwise stated) where 80% were excitatory (regular spiking) and 20% were inhibitory (fast spiking) cells. All dynamical equations were integrated using a forward Euler scheme with time step Δ*t* expressed in milliseconds. The membrane potential (*v*_*i*_) and recovery variable (*u*_*i*_) of each cell was modeled using the framework of Izhikevich (IZ) [37],

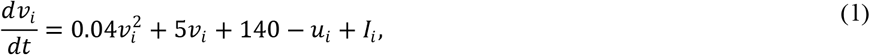

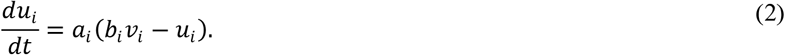

Spikes were triggered when *v*_*i*_>*v*_thresh_, where *v*_thresh_=30 mV. Following a spike, the membrane potential and recovery variable were reset to *v*_*i*_←*c*_*i*_ and *u*_*i*_←*u*_*i*_+*d*_*i*_, where *a*_*i*_, *b*_*i*_, *c*_*i*_, and *d*_*i*_ are parameters (set to *a*_*i*_=0.02, *b*_*i*_=0.2, *c*_*i*_=-65, and *d*_*i*_=8 by default). The synaptic current into each cell was

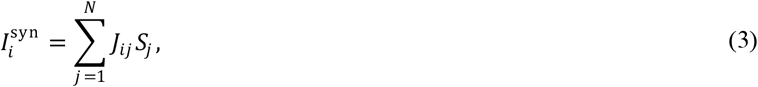

given synaptic weight *J*_*ij*_ from neuron *j* to *i*, and *S*_*j*_=1 if *v*_*j*_>*v*_thresh_, and *S*_*j*_=0 otherwise. For simplicity, synaptic currents are modeled as instantaneous contributions proportional to presynaptic spike events, without additional synaptic filtering. The total input current to neuron *i* is given by

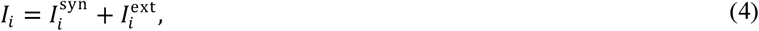

where 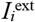represents external input originating from the neural controller, as described next.

### B. Closed-loop neural controller

The neural controller was designed as a reservoir network comprised of *N*=100 units that sends an activation signal to the simulated brain network. In turn, the brain network delivered activity back to the controller in real-time to guide the training process. Readout weights projecting from the reservoir network to the brain network were trained to perform a particular goal. Two goals were examined, where the controller either induced oscillations in the brain network or suppressed them. These two goals correspond to a broad range of applications of neural controllers, where a brain region is activated through an oscillatory input or where pathological oscillations are suppressed [12]. In the case where the controller’s goal was to induce oscillations, a sine wave was injected as input to the reservoir network which served as an oscillatory drive to train the controller. In this framework, we assume prior knowledge of the desired target stimulation to be injected in the simulated brain network. This corresponds to a scenario where a clinician has identified that a particular frequency of stimulation is beneficial to alleviate disease-related symptoms. The broader question of how to leverage machine learning approaches to identify optimal stimulation parameters based on individual patient symptoms is outside the scope of the current work.

Formally, given a reservoir state *x*(*t*)∈ℝ^Nres^, a recurrent weight matrix *J*∈ℝ^Nres×Nres^, an input weight vector *w*_in_∈ℝ^N^, an input signal *u*(*t*) designed as a 30 Hz sine wave, and a readout weight vector *w*_out_∈ℝ^N^, the reservoir dynamics are computed as

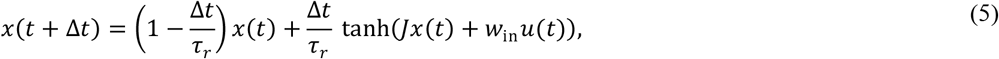

where τ_*r*_=10 ms. The reservoir output was given by

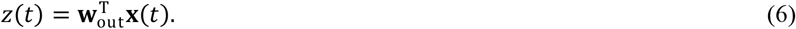

The scalar output *z*(*t*) is used solely for training the controller, while signals from the stimulating electrode are constructed directly from the reservoir state **x**(*t*). This output was converted to local field potentials (LFPs) before feeding to the spiking neurons, given that most measurements relating to diseases such as Parkinson’s disease come from LFP readings [38]. We assume a number of stimulating electrodes *k*=1,…,*N*_elect_ that represent a linear mixture of reservoir states,

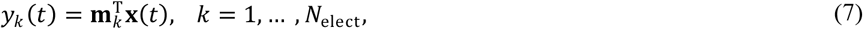

where **m**_*k*_∈ℝ^Nres^ is the spatial weight vector of electrode *k* and *N*_elect_ denotes the number of stimulating electrodes. Each stimulating electrode signal is low-pass filtered using a first-order exponential filter,

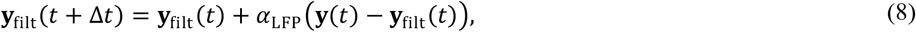

with

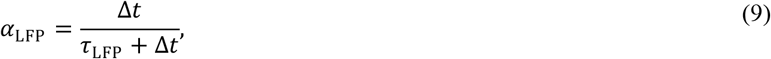

where τ_LFP_ is a low-pass time constant (in units of ms). Stimulating electrode signals are pooled to yield an LFP drive. In order to induce neuronal oscillations, we set

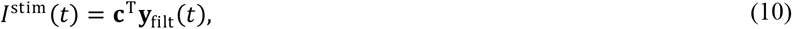

where **c**∈ℝ^Nelect^ is the electrode mixing vector using a uniform average. The injected current into each neuron is

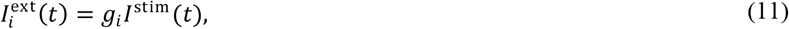

where *g*_i_=1 is an electrode-to-neuron coupling strength chosen uniformly across neurons.

The readout weights of the recurrent network were trained using CLEAR-FORCE. Unlike standard FORCE learning, where feedback is implicit and training is guided solely by output error, CLEAR-FORCE explicitly incorporates a second error term derived from the controlled spiking network itself. This hybrid error formulation allows the controller to adapt to non-stationary neural responses without requiring an explicit model of the target network.

First, reservoir error based on standard FORCE learning is computed as the difference between the network output *z*(*t*) and a desired target *z*\*(*t*),

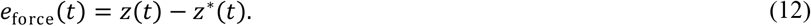

Second, IZ error was computed based on the difference between the average membrane potential 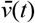 of the spiking network and the target *z*\*(*t*),

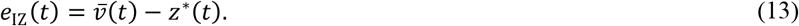

The use of the population-averaged membrane potential as a feedback signal serves two practical purposes. First, it provides a coarse-grained measure of network state that is closely related to local field potential (LFP) signals, which are commonly used in clinical settings. Second, it is computationally simple and hardware-efficient, making it well suited for real-time closed-loop implementations (see below).

Following equations (12-13), the learning process considered both the output of the stimulator and the response of the simulated brain network in order to adjust its weights. The two sources of error are combined into a total error *e*(*t*) as

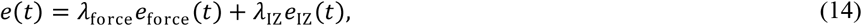

where fixed parameters λ_force_ and λ_IZ_ control the relative impact of each error source. When λ_IZ_=0, this is equivalent to standard FORCE learning used as an open-loop controller where brain signals are not fed back to the reservoir network.

Weight updates are performed using a recursive least squares approach by computing the projection

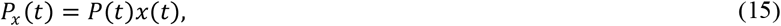

where *P*(*t*) is the running inverse of the correlation matrix obtained from recurrent network activity. The update to the readout weights is

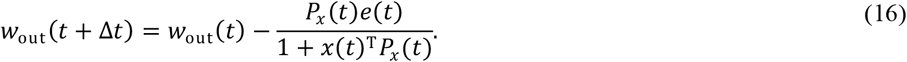

The update to the correlation matrix was

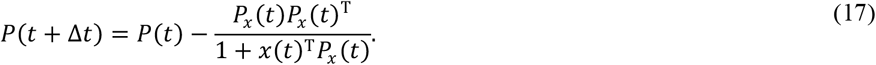

As with standard FORCE learning, stability is maintained through regularization of the correlation matrix and bounded reservoir activity. Next, we examined two scenarios: (1) where the neural controller generated oscillations in the simulated brain network; and (2) where the controller suppressed pathological oscillations.

## III. Inducing neural oscillations

Several clinical applications have considered a neural controller that induces neural activity in a target brain network. Broadly speaking, creating desirable patterns of brain activity may provide a way to restore or even enhance normal brain function. For instance, synchronous neural activity in the beta band (13-30 Hz) has been examined for controlling stimulation in Parkinson’s disease and essential tremor [4-6]. Further, gamma power (30-80 Hz) has been employed in the treatment of drug resistant depression [39-42] and to enhance reading performance in patients with dyslexia [43].

To induce oscillatory activity in the simulated brain network, we set the input to the reservoir network *u*(*t*) to a 30 Hz sine wave. We then began by examining baseline activity with the controller turned OFF. The simulated brain network showed spiking activity that was sparse and non-oscillatory (Fig. 3a). When turning the controller to ON, we saw the emergence of strong oscillations (Fig. 3b). While a spectrogram of neural activity showed no predominant band-specific power in the OFF mode (Fig. 3c), the ON mode showed heightened power at the target frequency (30 Hz) (Fig. 3d). Hence, the controller was effective at generating oscillatory activity in the simulated brain network.

**Fig. 3.**
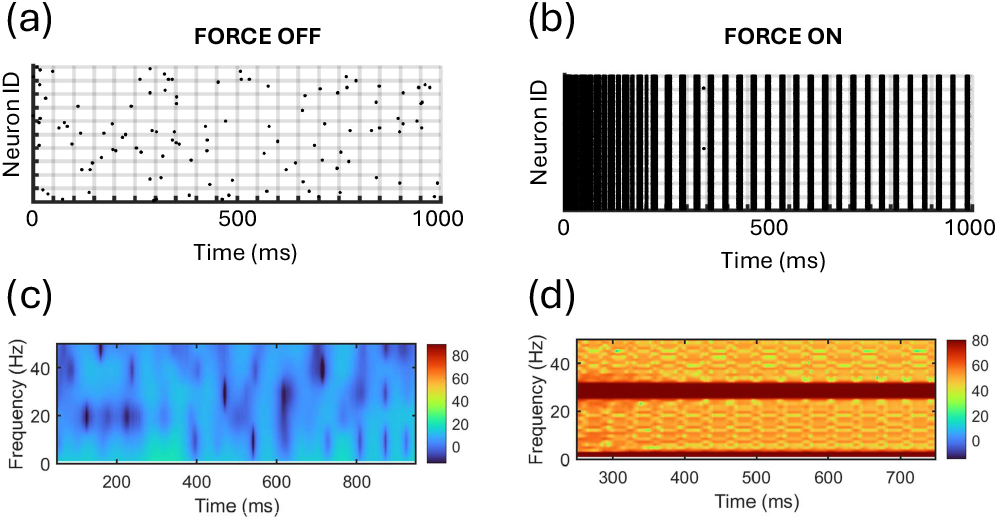
Inducing neural oscillations with CLEAR-FORCE. (a) Spike raster showing neural activity of the spiking network when the controller was OFF. (b) Turning ON the controller resulted in the emergence of sustained oscillations. (c) Spectrogram of neural activity with controller turned OFF. (d) Spectrograph with controller ON showing enhanced activity around 30 Hz.

Next, we examined a broad range of target oscillations, from 5-30 Hz, as well as a range of values of λ_IZ_ which determines the impact of feedback error from the brain network to the controller (see Equation (13)). Overall, we found a similar profile across oscillation frequencies: error was lowest when λ_IZ_ was set to a value above zero, showing that training the controller in a closed-loop fashion improved the ability of the system to induce brain oscillations compared to an open loop framework (λ_IZ_=0) (Fig. 4a). Across a range of physiologically-relevant stimulation frequencies, CLEAR-FORCE (λ_IZ_>0) outperformed an open-loop controller (λ_IZ_=0). The optimal value of λ_IZ_ varied between 3.1 and 4.7 (Table 1), where error is computed using Equation 14. It is interesting to note that a low value of λ_IZ_ resulted in higher error than λ_IZ_=0, suggesting that open loop stimulation outperformed a closed-loop controller when the feedback signal was too weak. Increasing λ_IZ_ to higher values was necessary for the closed-loop controller to surpass the open-loop performance.

**TABLE I.** Learning to synchronize a target network with CLEAR-FORCE.

| Stimulation Frequency (Hz) | Open-loop FORCE error | CLEAR-FORCE error | Optimal $\lambda_{IZ}$ |
| --- | --- | --- | --- |
| 30 | 15.147 (2.43) | 6.642 (0.07) | 4.1 |
| 25 | 14.516 (2.37) | 8.291 (0.09) | 4.0 |
| 20 | 14.257 (2.87) | 7.073 (0.07) | 3.3 |
| 15 | 13.899 (1.32) | 6.854 (0.06) | 3.6 |
| 12 | 13.488 (1.13) | 7.918 (0.06) | 3.1 |
| 10 | 12.883 (1.19) | 7.198 (0.09) | 3.4 |
| 5 | 12.668 (1.53) | 7.975 (0.09) | 4.7 |

Note. Values in table are means over 100 independent realizations. Numbers of parentheses are SEM.

**Fig. 4.**
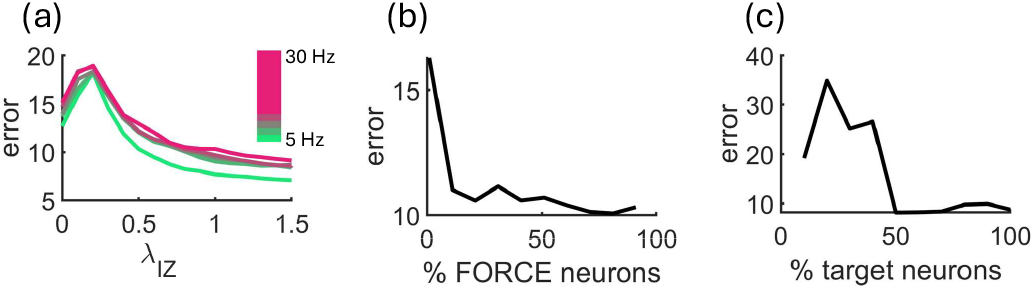
Impact of CLEAR-FORCE on generating neural oscillations in the simulated brain. (a) Controller error vs. strength of the closed-loop feedback (λ_IZ_). (b) Controller error vs. percentage of units in the reservoir that contributed to the output of the controller. (c) Controller error vs. percentage of spiking neurons targeted by stimulation.

In follow-up simulations, we altered the percentage of recurrent units in the CLEAR-FORCE network that contributed to the output of the controller (1-100%) and found that error decreased with a higher percentage of units. Error plateaued around 75 units (Fig. 4b). Hence, a small number of reservoir units were sufficient to induce oscillations in the simulated brain network. Similarly, we altered the number of spiking neurons from the simulated brain that were activated by the controller and found that approximately 50% of neurons were sufficient to induce low error (Fig. 4c). Hence, even partial activation of a brain network was enough to generate reliable oscillatory behavior.

Finally, we considered a scenario where the simulated brain network generated oscillations even when the controller was OFF (Fig. 5a). Here, the goal was to enhance existing oscillations. This was indeed found when switching the controller to ON (Fig. 5b). When altering the value of λ_IZ_, results were similar to those obtained with a non-oscillating brain network, namely that closed-loop stimulation (λ_IZ_∼1.5) was superior to an open-loop scenario (λ_IZ_=0) (Fig. 5c).

**Fig. 5.**
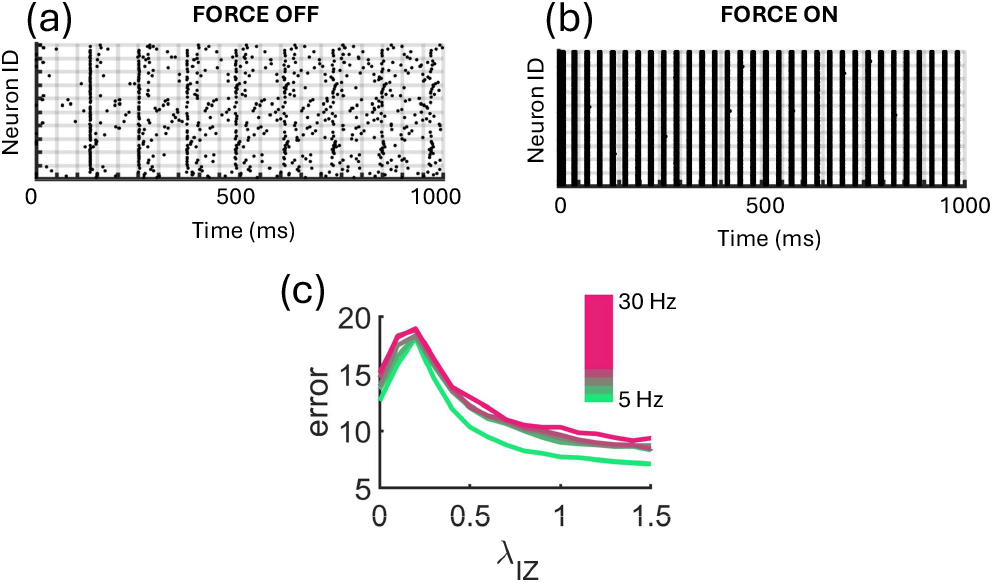
CLEAR-FORCE trained to modulate oscillations in a simulated brain network where baseline oscillations are already present. (a) Baseline oscillations in a spiking network with the controller switched OFF. (b) Sustained oscillations are produced when the controller is ON. (c) Controller error vs. strength of the closed-loop feedback (λ_IZ_).

In sum, a neural controller based on CLEAR-FORCE was effective at inducing brain oscillations in a simulated brain network composed of spiking neurons. Next, we considered whether the opposite scenario was also achievable: where the controller dampens a brain network that exhibits pathological oscillations.

## IV. Suppression of neural oscillations

Both theoretical [44] and experimental work [45-48] describe scenarios where pathological neural bursts and synchrony need to be removed. Closed-loop algorithms have been described as a way to desynchronize neuronal populations [49].

To train the neural controller to perform burst suppression, we replaced the external stimulation from Eq.10 to

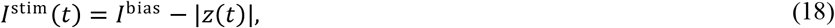

where *I*^bias^ is a tonic drive delivered to the simulated brain network. Subtracting the reservoir output *z*(*t*) from the tonic drive *I*^bias^ reduced the drive delivered to the spiking network, creating a form of suppression that dampens oscillatory bursts. Finally, to prevent oscillations induced by external input to the reservoir, we set *u*(*t*) to zero.

### A. Learning to suppress network bursts

To test the ability of the model to perform burst suppression, we simulated a spiking network with strong tonic drive (*I*^bias^=5, in arbitrary units) and trained a neural controller using a reservoir network with CLEAR-FORCE. The closed-loop controller selectively quenched transient, high-synchrony events while preserving low-rate, unsynchronized background activity (Fig 6). While this is was a simple task, as the reservoir network simply had to increase its tonic activity in order to suppress bursts, a more complex task involving real-time burst suppression is presented in the next section.

**Fig. 6.**
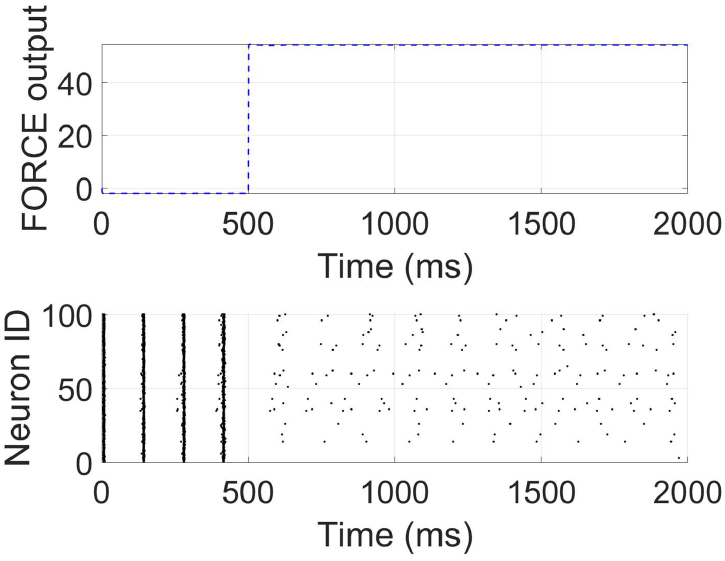
Suppression of burst activity with CLEAR-FORCE while maintaining network activity that is not globally synchronized.

### B. Real-time burst suppression

A series of simulations were employed to show how CLEAR-FORCE can learn in real time to suppress a burst that emerges sporadically during spiking activity. Leaving the controller OFF, we induced a burst in the spiking network by increasing the external drive during a delimited time window (5000 – 6000 ms). In the same simulation, but with the controller ON, the same drive did not result in a burst (Fig. 7). Thus, the controller successfully suppressed a burst that emerged in real time during spiking dynamics.

**Fig. 7.**
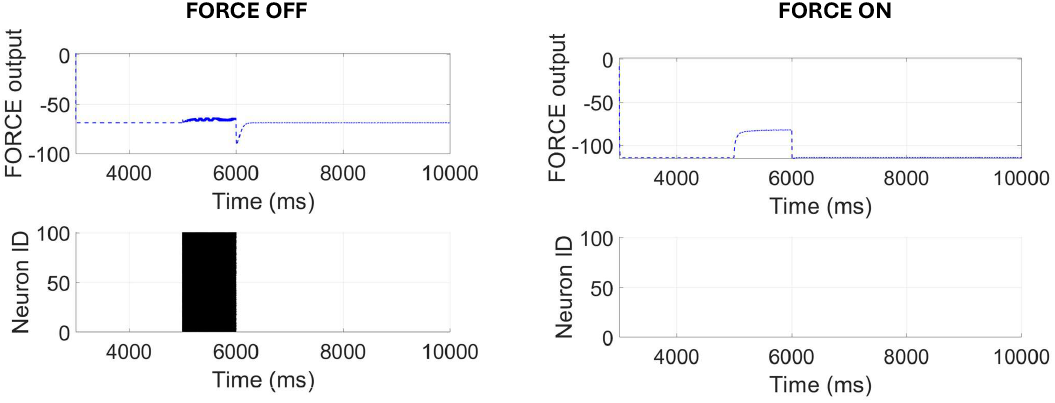
Real-time burst suppression with the CLEAR-FORCE controller.

Next, we examined scenarios where a controller targeted specific neuronal subtypes including excitatory (E) and inhibitory (I) spiking neurons. Within the spiking network, we also manipulated which neurons generated the bursts (E vs. I neurons). We examined the firing rate of the spiking neurons to see whether CLEAR-FORCE was successful at suppressing bursts.

A summary of results is presented in Table II. Overall, CLEAR-FORCE was successful at suppressing bursting activity, as the resulting firing rates were low for both E and I neurons (<2 Hz) regardless of the burst origin or the neuronal subtype targeted by the controller. By comparison, firing rates were consistently higher when the controller was OFF. Hence, burst suppression by CLEAR-FORCE was possible regardless of whether bursts originated from E or I neurons and whether the controller acted primarily on E, I, or both subtypes of cells.

**TABLE II.** Neuron-specific Burst suppression with CLEAR-FORCE.

| *Burst origin* | *CLEAR-FORCE target* |  |  |
| --- | --- | --- | --- |
|  | *I neurons* | *E neurons* | *E and I neurons* |
| *I neurons* | ON: 1.44 Hz (0.003)<br>OFF: 7.28 (0.025) | ON: 1.44 Hz (0.003)<br>OFF: 7.23 Hz (0.006) | ON: 1.44 Hz (0.065)<br>OFF: 7.27 Hz (0.0013) |
| *E neurons* | ON: 1.44 Hz (0.003)<br>OFF: 7.21 Hz (0.025) | ON: 1.44 Hz (0.003)<br>OFF: 7.21 Hz (0.005) | ON: 1.44 Hz (0.055)<br>OFF: 7.27 Hz (0.0011) |
| *E + I neurons* | ON: 1.35 Hz (0.003)<br>OFF: 7.22 Hz (0.025) | ON: 1.44 Hz (0.003)<br>OFF: 7.21 Hz (0.006) | ON: 1.53 Hz (0.048)<br>OFF: 7.20 Hz (0.009) |

## V. NEUROMORPHIC POWER CONSUMPTION

Finally, we aimed to estimate the power consumption of the reservoir network for implementation in a neuromorphic controller. Simulations were performed where we varied the number of spiking neurons (*N*_spk_) while keeping the number of reservoir units (*N*_res_) fixed at 1,000. In this case, the controller error increased approximately linearly, with a linear fit explaining nearly all variance (coefficient of determination: R^2^=0.9965) and an exponential fit performing poorly (R^2^=0.1829) (Fig. 8a). Conversely, when varying the number of reservoir units and fixing *N*_spk_ at 1,000, error decreased exponentially, with an exponential model providing an excellent fit (R^2^=0.9942) compared to a much weaker linear fit (R^2^=0.6823) (Fig. 8b).

**Fig. 8.**
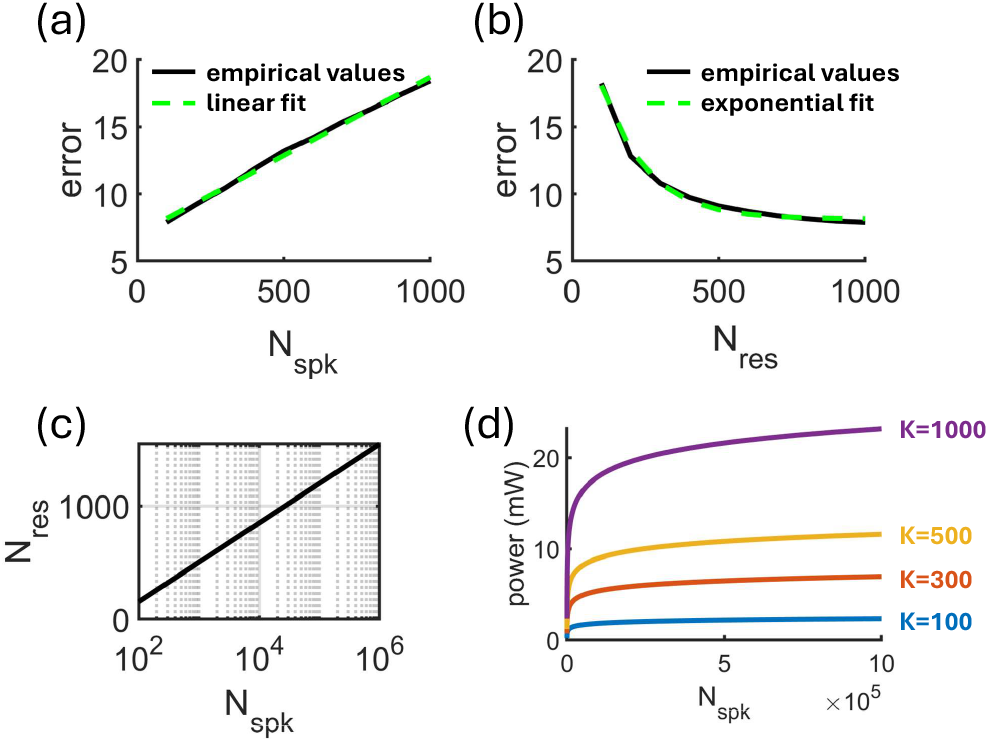
Estimating CLEAR-FORCE power consumption for neuromorphic applications of brain stimulation. (a) Linear increase in error as the number of spiking neurons (*N*_spk_) increases. (b) Exponential decrease in error as the number of reservoir neurons (*N*_res_) increases. (c) Logarithmic relation between *N*_spk_ and *N*_res_. (d) Estimated dynamic compute power relative to *N*_spk_ and *K* (average number of connections per reservoir unit).

These results show that scaling to a larger spiking network increases error linearly, while increasing the reservoir size decreases error exponentially. Thus, a logarithmic relation characterizes the relation between *N*_res_ and *N*_spk_ (Fig. 8c):

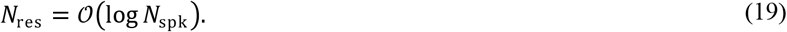

Using the parameters obtained from the best-fitting linear and exponential functions, a controller with roughly *N*_res_=1,500 to 2,000 units would be of the correct order of magnitude to interact with large subcortical populations such as the human striatum, where medium spiny neurons are on the order of ∼10^8^ neurons [50].

To estimate the power consumption associated with a reservoir of that size, we assumed that the dynamic power of a neuromorphic reservoir network scales approximately linearly with the number of reservoir units *N*_res_. To simplify calculations, we accounted only for controller computation; in a closed-loop system, additional power is required for sensing, stimulating output, and communication, which are not included in this calculation. The main contribution of power consumption arises from synaptic operations, which scale as

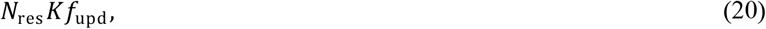

multiplied by a given energy cost per operation, where *K* denotes the average number of connections per reservoir unit, and *f*_upd_ denotes the update rate of the reservoir dynamics (in Hz). Using representative values (*N*_res_≈2,000, *K*∼10^2^ – 10^3^, and *f*_upd_∼10^3^ Hz, with tens of pJ per synaptic operation), this yields an estimated dynamic computing power in the range of a few to a few tens of milliwatts (Fig. 8d). These power levels are compatible with low-milliwatt budgets reported for neuromorphic hardware platforms, supporting the feasibility of portable reservoir-based neural controllers [51].

## VI. Discussion

The present work introduced CLEAR-FORCE as a reservoir-based closed-loop controller capable of both inducing and suppressing neural oscillations in a spiking network model. A central contribution of this framework is that adaptive control is achieved by training a reservoir network which is relevant for real-time neuromodulation, where power consumption and scalability are important constraints [52]. Because small recurrent networks are able to control complex spiking networks that evolve over time, they are well suited as adaptive neural controllers that maintain low energy costs.

From a reservoir computing standpoint, the central contribution of CLEAR-FORCE resides in its learning rule, which jointly optimizes two error signals: a conventional reservoir output error and a feedback error derived from the dynamics of the controlled system. In this way, CLEAR-FORCE extends standard FORCE learning from a simple pattern matching procedure to a closed-loop optimization framework. This dual-error formulation allows the reservoir readout to adapt not only to a prescribed target signal but also to deviations in the system being controlled, enabling robust performance under non-stationary and partially unpredictable dynamics.

Related approaches based on high-performance brain-computer interfaces can decode motor and cognitive signals from large populations of neurons [25,27]. While these systems often rely on offline or power-intensive architectures, CLEAR-FORCE highlights how closed-loop control can be achieved using an energy efficient architecture, hence opening a pathway to deployable hardware for clinical applications. This aligns with recent work which demonstrates efficient neuromorphic implementations of reservoir computing [51].

In contrast to time-adaptive Bayesian optimization methods that use probabilistic surrogate modeling and iterative parameter updates [19], our reservoir controller with FORCE learning offers a model-free approach capable of continuous, real-time adaptation. By directly learning the nonlinear mappings between neural activity and stimulation, it enables more flexible control in complex neural environments.

Some limitations of the present work point to directions for future research. First, in simulations where the controller induced oscillations, the target of the reservoir remained fixed. In clinical settings, however, stimulation would have to adapt over time based on symptoms and changes in neural states. This type of dynamic adaptation could be considered in extensions of CLEAR-FORCE.

Second, the spiking network explored here uses a single Izhikevich parameter regime. Interestingly, varying between regular-spiking and bursting regimes could serve as a gateway to personalized neural models that better approximate patient-specific dynamics. Such personalized models could be used *in silico* to test stimulation strategies prior to clinical deployment, potentially reducing trial-and-error during parameter tuning.

Third, the reservoir network is not constrained in a way that would provide the minimum effective stimulation to either induce or suppress brain oscillations. This is particularly relevant to clinical applications where transcranial stimulation may be accompanied by negative side-effects. This issue could be addressed by adding an explicit term to the error function in order to constrain the reservoir to a solution with a low amplitude of stimulation.

An especially promising extension would be to incorporate synaptic plasticity within the spiking network itself. Including mechanisms such as spike-timing-dependent plasticity (STDP) would modulate long-term network reorganization. In this scenario, CLEAR-FORCE could apply stimulation in order to induce plasticity with the goal of eliminating network regimes that give rise to pathological oscillations, capturing experimentally observed effects of stimulation on brain plasticity [53,54]. Finding the optimal stimulation parameters to induce brain plasticity remains an open theoretical question and represents a fertile area for future work.

Finally, while the present results demonstrate that CLEAR-FORCE can suppress network bursts regardless of whether excitatory or inhibitory neurons are targeted, a more refined controller could employ separate objective functions for excitatory and inhibitory populations. However, implementing this strategy in hardware would require reliable online identification of excitatory and inhibitory neurons from extracellular spike data, a challenging problem that we have begun to address with unsupervised processing pipelines in high-density recordings [55].

In conclusion, this work positions CLEAR-FORCE as a promising computational framework for adaptive, low-power neural control. By leveraging the dynamics of reservoir computing, this approach provides a first step towards bridging advances in reservoir networks with practical constraints of implantable neuromodulation hardware. As neural interfaces continue to scale in throughput, reservoir-based controllers offer a compelling route toward personalized, energy-efficient, and clinically viable brain stimulation technologies.

## ACKNOWLEDGMENT

This work was funded by a grant to J.P.T. from the Natural Sciences and Engineering Research Council of Canada (NSERC Grant No. 210977). Authors are thankful for discussions with the laboratory of Christopher Pack (McGill University).

